# 3D Hyaluronic Acid Hydrogels for Modeling Oligodendrocyte Progenitor Cell Behavior as a Function of Matrix Stiffness

**DOI:** 10.1101/2020.04.01.020412

**Authors:** Deniz B. Unal, Steven R. Caliari, Kyle J. Lampe

## Abstract

The lack of regenerative solutions for demyelination within the central nervous system (CNS) motivates the need for better understanding of the oligodendrocytes that give rise to myelination. In this work, we introduce a 3D hyaluronic acid (HA) hydrogel system to study the effects of mechanical properties on the behavior of oligodendrocyte progenitor cells (OPCs), the cells that differentiate into myelin-producing oligodendrocytes in the CNS. We tuned the stiffness of the hydrogels to match brain tissue (storage modulus 200 – 2000 Pa) and studied the effects of stiffness on metabolic activity, proliferation, and cell morphology of OPCs over a 7 day period. Although hydrogel mesh size decreased with increasing stiffness, all hydrogel groups facilitated OPC proliferation and mitochondrial metabolic activity to similar degrees. However, OPCs in the two lower stiffness hydrogel groups (169.8 ± 42.1 Pa and 793.9 ± 203.3 Pa) supported greater adenosine triphosphate (ATP) levels per cell than the highest stiffness hydrogels (2178.7 ± 127.2 Pa). Lower stiffness hydrogels also supported higher levels of cell viability and larger cell spheroid formation compared to the highest stiffness hydrogels. Together, these data suggest that 3D HA hydrogels are a useful platform for studying OPC behavior and that OPC growth/metabolic health may be favored in lower stiffness microenvironments mimicking brain tissue mechanics.

## Introduction

One of the major life-altering effects of central nervous system (CNS) disorders is demyelination, with multiple sclerosis being the most common disease pathology.^1^ Without the lipid-based myelin sheath to insulate neuronal axons, these axons that rely on efficient node to node conduction revert to inefficient continuous conduction. In turn, this leads to protease activation, axonal degradation, axonal loss, and ultimately neuronal death.^2^ This motivates the development of strategies to promote “remyelination” or the regeneration of the myelin sheath after degeneration. Oligodendrocyte progenitor cells (OPCs) of the central nervous system are one of the most important cell types in this research.^3^ OPCs are distributed through the white matter in the adult brain and differentiate into mature oligodendrocytes.^4^ Oligodendrocytes myelinate the axons of neurons in the CNS (counterparts to Schwann cells in the peripheral nervous system) to increase efficiency and precision of electrical impulse conduction.^5^ Known as the recapitulation hypothesis, remyelination after injury may follow similar processes to developmental myelination. In such a scenario, OPC development and maturation could be leveraged in bioengineering strategies to treat multiple sclerosis and other demyelinating diseases.^6^

Hydrogels are useful biomaterials for modeling CNS extracellular environments as they are mechanically compliant, permit biomolecule transport, and facilitate cell behaviors including adhesion, proliferation, and migration.^7–9^ OPCs and their precursors (neural stem cells) have been studied in polyethylene glycol (PEG) hydrogels^10^ as well as collagen, agarose, chitosan, and fibrin hydrogels.^11,12^ Hyaluronic acid (HA) is an excellent candidate for engineering brain-like extracellular matrix (ECM) microenvironments^13,14^ due to its abundance in the CNS^15–17^ and its anti-inflammatory role in brain and spinal cord injuries.^18^

Hydrogel cell culture models enable interrogation of the roles that extracellular cues such as matrix stiffness play in regulating salient cell behaviors.^10,19–21^ Jagielska et al. showed that OPCs are mechanosensitive, with proliferation being greatest on 2D substrates with Young’s elastic moduli of 700 Pa.^22^ 3D culture systems offer the potential to study cell-cell and cell-matrix interactions as well as biotransport conditions in a more physiologically-relevant setting. We previously showed that OPC metabolic activity was greatest within 3D PEG hydrogels with storage moduli of 240 ± 84 Pa in comparison to 630 ± 79 Pa.^23^ In this work, we used 3D HA hydrogels to interrogate the effects of stiffness and mesh size on OPC proliferation, metabolic activity, and spheroid volume.

## Materials & Methods

### Cell culture

Green fluorescent protein positive (GFP+) mosaic analysis with double markers (MADM) OPCs were expanded on T75 (Corning CellBind) treated tissue culture plates. MADM cells are derived from mouse glioma cells that express many OPC markers.^23^ Tissue-cultured plates were coated with polyornithine (10 µg/mL) and rinsed thrice with Dulbecco’s phosphate buffered saline (PBS). OPC proliferation media consisted of Dulbecco’s modified Eagle’s medium (DMEM, Life Technologies) with 4 mM L-glutamine and 1 mM sodium pyruvate (Life Technologies), N2 supplement (Life Technologies) and B27 supplement (Life Technologies), and 1% penicillin-streptomycin (Life Technologies). Cells were seeded at 0.5 x 10^4^ cells/cm^2^ in 8 mL of proliferation media. Media was changed every two days. Passaging was performed when cells reached 80% confluence. Cyropreserved cells were passaged at least once before use in hydrogel experiments. Cells from passage numbers 10-25 were used.

### NorHA macromer synthesis

#### HA-TBA synthesis

Research grade sodium hyaluronate (Lifecore Biomedical, 61.8 kDa) was converted to its tetrabutylammonium (TBA) salt (HA-TBA) to prepare for dissolution in DMSO.^24^ Sodium hyaluronate was dissolved in DI water at 2 wt% and Dowex resin (50W x 200) was added at a ratio of 3 g resin/1 g HA.^24^ The resin exchanges sodium ions for hydrogen ions, making the solution strongly acidic. The solution was stirred for ∼ 2 hrs. Stirring was stopped to allow the resin to settle to the bottom of the flask. The solution was then vacuum filtered until the solution was clear to remove all resin. The solution was titrated to pH 7.02-7.05 with TBA-OH. The HA-TBA solution was then frozen at -80 °C and lyophilized. Tubes were purged with N2 and stored at -20 °C.

#### NorHA synthesis

In a dried and stoppered round bottom flask, HA-TBA and Nor-amine (5- norbornene-2-methylamine) were added. Nor-amine was added to modify ∼ 20-30% of HA disaccharides. Anhydrous DMSO was added via cannulation to the flask (∼ 5 mL per 0.1 g). Once the HA-TBA was fully dissolved, benzotriazol-1-yloxytris(dimethylamino)phosphonium hexafluorophosphate (BOP) was added at a 0.3:1 molar ratio of BOP to TBA functional groups on the HA-TBA via cannulation with the BOP dissolved in ∼ 20 mL of anhydrous DMSO. The reaction was allowed to proceed for ∼ 2 h and then quenched with ∼ 10 mL of DI water. The solution was transferred to dialysis tubing (molecular weight cutoff: 6-8 kDa) and put on dialysis for 5 days, the first 3 of which included ∼ 5 g of NaCl in ∼ 5 gal of DI water to help remove TBA. Through filtration, the side-products from the BOP coupling were removed. The solution was then returned to dialysis for 3-5 days. Finally, the solution was transferred to 50 mL tubes, frozen at - 80 °C, and lyophilized to dry. A representative ^1^H NMR spectrum can be found in **Figure S1**.

#### Hydrogel preparation

Hydrogel precursor solutions were prepared to final concentrations of 1–2 wt% NorHA macromer, 0.0328 wt% lithium phenyl-2,4,6-trimethylbenzoylphosphonate (LAP) photoinitiator, and dithiothreitol (DTT, Sigma-Aldrich) crosslinker at a 0.315:1 ratio of thiol:norbornene in PBS. Solutions were sterilized via germicidal UV irradiation for 2.5 h. OPC MADM cells were combined with the precursor solution and encapsulated in hydrogels at a final concentration of 5 × 10^6^ cells/mL using 365 nm UV light (4 mW/cm^2^, 2 min).

#### Oscillatory shear rheology

Oscillatory shear rheology was used to assess gelation kinetics and hydrogel viscoelastic properties. 50 µL of hydrogel precursor solution was pipetted onto a UV-configured plate of an Anton Paar MCR 302 rheometer. A 25 mm diameter smooth cone and plate with a 0.505° angle was used for all studies. Hydrogels were cured *in situ* at 0.1% oscillatory strain, 10 rad/s oscillation frequency, and 4 mW/cm^2^ UV light intensity (365 nm). After 30 s of pure oscillatory motion, light exposure was introduced underneath the plate for 120 s. Time sweep data was recorded for an additional 60 s after light exposure ceased.

#### Swelling studies

360 µL hydrogels were fabricated in 3 mL syringes using the formulations and polymerization conditions already described. Samples were then soaked in PBS at 37 °C. At time points of 0, 24, and 48 h after gelation, samples were weighed after blotting away excess liquid. These samples were then lyophilized and the dry mass was recorded. Two experimental replicates were conducted with *n* = 3-4 per sample type per time point. Flory-Rehner calculations^25^ were used to determine mass and volumetric swelling ratios (**Table S1**), molecular weight between crosslinks, and mesh size. First the molecular weight between crosslinks was determined through the equilibrium swelling theory for crosslinked polymers^26^:

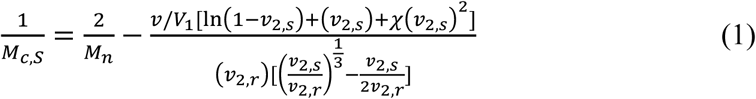

Where M_c,S_ is the molecular weight between crosslinks estimated from swelling (g/mol), Mn is the molecular weight of the macromer (61,800 g/mol), *v* is the specific volume of the macromer (0.8137 mL/g), *V*_1_ is the molar volume of the solvent (18 mol/cm^3^), χ is th e polymer solvent interaction parameter, *v*_2,*r*_ is the volume fraction of polymer in the relaxed state, and *v*_2,*s*_ is the volume fraction of polymer in the swollen state (see supplemental information for more details). The polymer solvent interaction parameter can be taken as 0.473 because of the similarities between the polysaccharides hyaluronic acid and dextran.^25,27^ The effective crosslinking density and mesh size calculations are then adapted from Leach et al.^27^

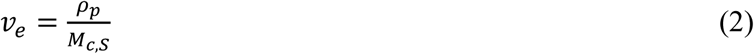

Where *v*_*e*_ is the effective crosslinking density (mol/cm^3^), and ρ_*p*_ is the density of the dry polymer (g/mL). This allows calculation of the mesh size using the equation below:

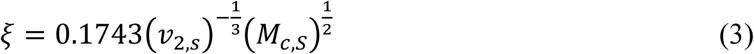

Where *ξ* is the mesh size (nm).

### Quantification of OPC mitochondrial metabolic activity

OPC mitochondrial metabolic activity was measured using the AlamarBlue assay. In AlamarBlue, the chemical compound resazurin is reduced to resorufin through cellular respiration. The metabolic activity is represented by percent reduction and can be measured through absorbance or fluorescence intensity. The Beer-Lambert law is used when measuring absorbance values. Equations 4, 5, and 6 describe the phenomena.

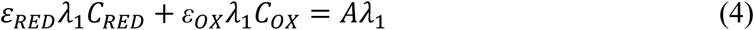

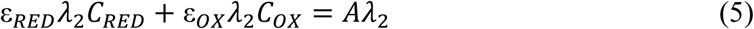

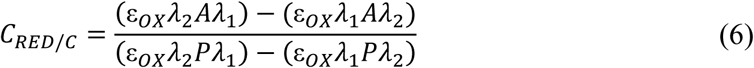

Where *ελ* is the molar extinction coefficient (oxidized or reduced form) at the excitation or emission wavelength (570 nm = *λ*_1_, 600 nm = *λ*_2_), *Aλ* is the absorbance value of the sample at the excitation or emission wavelength, *C* is the concentration of AlamarBlue (oxidized or reduced form), and *Pλ* is the absorbance of the control wells (excitation or emission). AlamarBlue reagent (ThermoFisher) was mixed with OPC proliferation media at a 1:9 ratio as specified by the manufacturer. Each hydrogel with encapsulated cells was incubated in 400 µL of this pre-made solution for 4 h at 37 °C. Hydrogels were then removed from solution and returned to OPC proliferation media for further culture. Remaining solution for each sample was pipetted into three 100 µL replicates along with three 100 µL replicates of blank AlamarBlue/media solution. Absorbance was read in a BMG Clariostar monochromator microplate reader.

### Live/Dead assay

OPC viability in HA hydrogels was measured using the Live/Dead assay. Each hydrogel was suspended in 1 mL of PBS plus glucose (PBSG) with the same concentration of glucose as in OPC proliferation media (25 mM). Live/Dead staining consisted of 4 µM ethidium homodimer (to label dead cells) with incubation at 37 °C for 60 minutes followed by rinsing 2x with PBSG. Live cells were visualized using GFP signal from MADM OPCs. Live/Dead images were collected using a Zeiss LSM 510 confocal microscope with ∼ 500 µm z-stacks (10 µm slices).

### Cell Glo

OPC ATP activity was measured using the Cell Glo assay. Hydrogel samples were homogenized in 400 uL of lysis buffer (20 mM Tris-HCl, 150 mM NaCl, 1 mM Na2EDTA, 1 mM EGTA, 1% Triton X-100, 2.5 mM sodium pyrophosphate, 1 mM β-glycerophosphate, 1 mM Na3VO4, 1 µg/mL leupeptin). First, a standard curve for ATP was made using the manufacturer’s protocol. Cell Glo reagent was prepared by adding 10 mL of CellTiter-Glo buffer to the CellTiter-Glo substrate to reconstitute the lyophilized enzyme mixture. Solution was gently mixed through inversion. Once completely dissolved, the solution was stored at – 20 °C in 1 mL aliquots. A white 384-well plate was used for the assay. 25 µL of each standard was added with three pipetting replicates. 5 µL of sample was diluted with 20 µL of PBS in each well. To all wells (standards and samples), 25 µL of Cell Glo reagent was added. The plate was incubated at room temperature for 10 minutes and luminescence was measured using a BMG Clariostar microplate reader. A standard curve was plotted relating luminescence values to ATP concentrations. Sample ATP concentrations were then determined based on the standard curve.

### Pico Green

OPC DNA content was measured using the Pico Green assay. Hydrogel samples were homogenized in 400 uL of lysis buffer (20 mM Tris-HCl, 150 mM NaCl, 1 mM Na2EDTA, 1 mM EGTA, 1% Triton X-100, 2.5 mM sodium pyrophosphate, 1 mM β-glycerophosphate, 1 mM Na3VO4, 1 µg/mL leupeptin). First, a standard curve for DNA was made using the manufacturer’s protocol. A black 384-well plate was used for the assay. Standards were pipetted at 20 µL with three pipetting replicates. Samples were pipetted with three pipetting replicates and three sample replicates at 5 µL sample volume and 15 µL 1x TE buffer (10 mM Tris-HCl, 1 mM disodium ethylenediaminetetraacetic acid (EDTA), pH 8.0). 20 µL of 1X Pico Green reagent was added to all the standards and samples. Fluorescence was measured in a BMG Clariostar microplate reader according to the manufacturer’s directions. A standard curve was plotted relating fluorescence values to DNA concentrations. Sample DNA concentrations were then determined based on the standard curve.

### Spheroid volume measurement

OPC spheroid volumes were measured using Live/Dead confocal images. Z-stacks of ∼ 500–1500 µm thickness (10 µm slices) were loaded in ImageJ. GFP and ethidium homodimer channel images were analyzed separately using the “3D object counter” module in ImageJ. This module performs image thresholding followed by counting and sizing of each spheroid. The volume of each GFP+ spheroid was determined. The average spheroid volume was calculated for each experimental condition and time point. In addition, the average spheroid volume was normalized to the hydrogel volume. The average hydrogel volume occupied by cell spheroids (a fraction of the hydrogel occupied by live cells) was calculated by summing the volume of spheroids within each image stack, normalizing to the hydrogel volume, and then averaging between three hydrogels per experimental condition per time point. A total of 3613–14870 spheroids were measured per experimental condition at a given time point. Three confocal image stacks were taken per sample. Two or three samples were analyzed from each experiment and time point. Five experimental replicates were performed, yielding 12-15 samples per condition per time point for each hydrogel experimental group.

### Statistical analysis

ATP, DNA, metabolic activity, and spheroid volume data were analyzed using two-way ANOVA followed by the Tukey post-hoc test with α-value of 0.05. Calculations were performed using SPSS software. Box plots cover the second and third data quartiles with error bars covering the first and fourth quartiles. Box plots also include marks for mean (*x*) and median (*bar*) data values.

## Results & Discussion

### Mechanical Properties of NorHA Hydrogels

The NorHA hydrogels were formed through the crosslinking of norbornene groups with dithiol crosslinker (DTT) via radical-mediated thiol-ene addition using a LAP photoinitiator that absorbs strongly at 365 nm (**Figure 1**). Norbornene groups preferably crosslink with thiyl radicals instead of with themselves, creating a phototunable hydrogel system where the degree of crosslinking is controlled by the molar ratio of thiol to norbornene groups.^24^ Gramlich et al. demonstrated the positive correlation between NorHA hydrogel stiffness and thiol to norbornene crosslinking ratio up to 0.8.^24^ We also previously demonstrated the suitability of NorHA hydrogels for 3D cell culture across a range of Young’s moduli from ∼ 1-15 kPa.^28^ In this work, the molar ratio of thiol to norbornene groups was kept constant at 0.315 while the NorHA macromer concentration was varied between 1 and 2 wt% to enable fabrication of hydrogels spanning a range of CNS-relevant stiffnesses.

**Figure 1:**
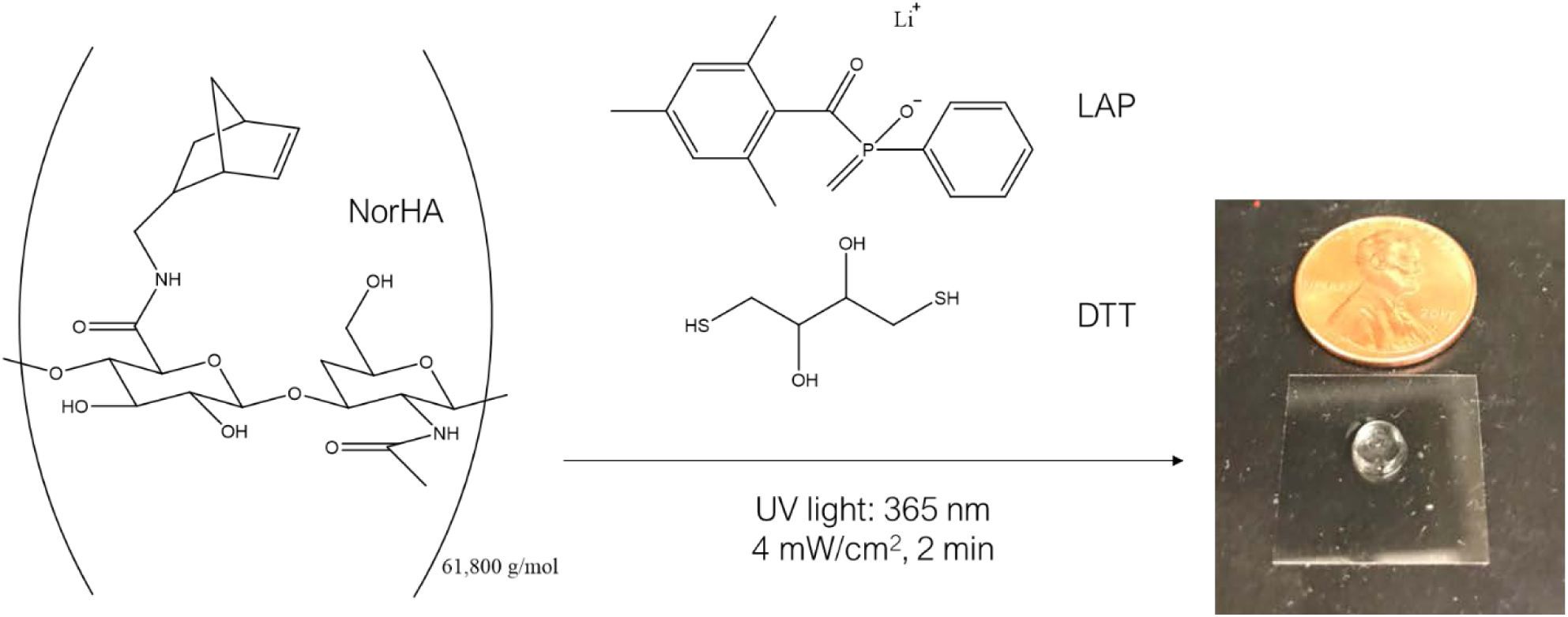
Schematic of hydrogel formation. 20-28% of the hyaluronic acid disaccharide repeat units were modified with norbornene functional groups. Norbornenes were crosslinked with dithiol crosslinker (DTT) to form hydrogels. Hydrogels were photocrosslinked under 4 mW/cm^2^ (365 nm) light for 2 minutes using LAP as the photoinitiator.

Cells sense mechanical cues and respond to them through mechanotransduction-mediated changes in morphology, metabolic activity, and proliferation.^29^ Storage modulus and mesh size are key properties affecting these cell traits.^23^ Storage modulus dictates the stiffness of the material while mesh size dictates the spacing between crosslinks and availability for cell colony growth and biomolecule diffusion. Storage modulus can be measured through oscillatory shear rheology (**Figure 2**).

**Figure 2:**
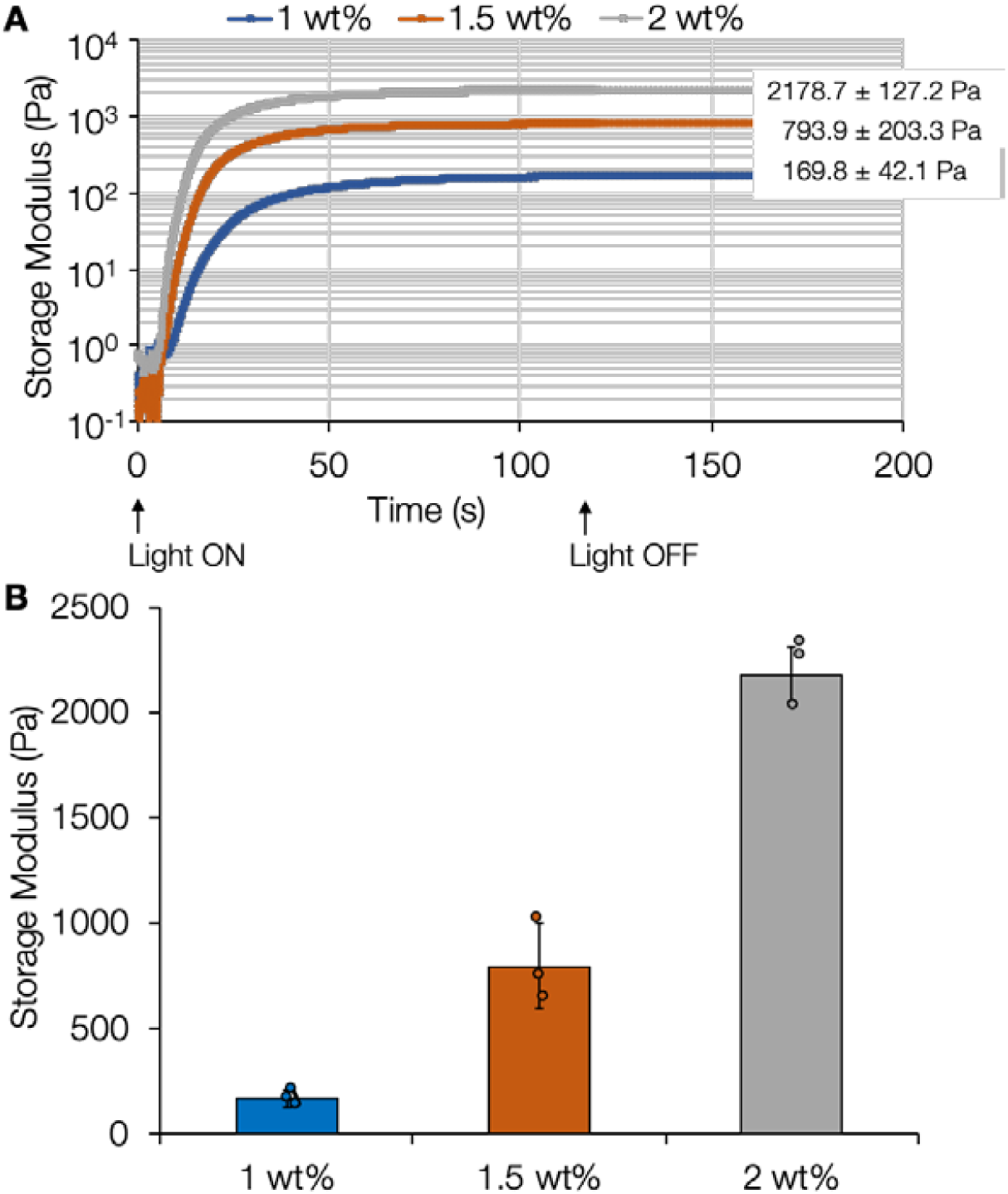
Rheological properties of crosslinked HA hydrogels. (A) Average storage modulus was determined from oscillatory shear rheology time sweeps performed on NorHA hydrogels crosslinked *in situ* at 1 wt% (wt/v), 1.5 wt%, or 2 wt%. Hydrogels were formed by exposure to UV light from 0 to 120 s. (B) Averaged plateau modulus (*bar*) was measured from individual trials (*circles*). Plateau storage modulus was calculated from the average modulus of 150–180 s (30-60 s after the light is turned off) for each trial. Loss moduli (∼ 1 Pa) do not significantly change between the experimental groups for these elastic hydrogels. *Error bars*: standard deviation, *n* = 3 hydrogels per experimental group.

Formulations of NorHA hydrogels were adjusted to match the range of brain tissue stiffness (i.e., storage moduli of 200-2000 Pa)^30^ with 1, 1.5, and 2 wt% formulations corresponding to storage moduli of 169.8 ± 42.1 Pa, 793.9 ± 203.3 Pa, and 2178 ± 127.2 Pa respectively. Hydrogels reached swelling equilibrium within 24 hours (**Table S1**). Mesh sizes at equilibrium were determined through Flory-Rehner polymer theory (Materials and Methods). In comparison to previous studies with PEG-dimethacrylate (PEGDM) hydrogels, the NorHA hydrogels have 6- 7 fold larger mesh sizes (**Table 1**) and much greater swelling ratios (**Table S1**), suggesting that the NorHA hydrogels could allow for greater cell growth or process extension while still appropriately mimicking CNS tissue stiffness. In previous work, the degree of methacrylation modification of PEGDM hydrogels was 82-86%^23^ while here the degree of modification of NorHA hydrogels was 20-28%. Leach et al. performed swelling studies with hydrogels made from glycidyl methacrylate-HA (GMHA) that was two orders of magnitude greater in molecular weight than the HA used here but only 5-11% functionalized with methacrylate groups.^27^ Molecular weight between crosslinks was also found to be two orders of magnitude greater (not reported here) compared to our NorHA hydrogels.^27^ Consequently, mesh sizes were an order of magnitude greater than those calculated for our NorHA hydrogels. Stiffnesses were comparable to NorHA and PEGDM hydrogels. While stiffness is typically inversely correlated with mesh size, stiffness is not a direct indication of mesh size. The results of our study and others^23,27^ show that hydrogel mesh size depends on a combination of the degree of functionalization, the molecular weight of the polymer, and the crosslinking density. Interestingly, these NorHA hydrogels, while exhibiting high swelling ratios, underwent relatively small increases in volume of 10-27% at 24 hours (**Table S1**). This may be advantageous in maintaining a high cell density, and thus maximizing important pericellular signaling.

**Table 1:** Hydrogel mesh sizes. Comparison of NorHA hydrogel mesh sizes with PEGDM hydrogel mesh sizes^23^ and GMHA hydrogel mesh sizes^27^ (* indicates complex shear modulus). Mesh sizes were calculated from swelling studies measuring equilibrium (swollen) wet mass and lyophilized dry mass to obtain mass / volumetric swelling ratios. *n* = 8 hydrogels per experimental group.

| Work Source | Molecular Weight (g/mol) | Degree of functionalization (%) | Sample Type | Storage Modulus (Pa) | Mesh Size (nm) |
| --- | --- | --- | --- | --- | --- |
| Present Study | 61,800 | 20-28 | 1 wt% NorHA | 169.8 ± 42.1 | 85.6 ± 2.6 |
| Present Study | 61,800 | 20-28 | 1.5 wt% NorHA | 793.9 ± 203.3 | 75.9 ± 3.5 |
| Present Study | 61,800 | 20-28 | 2 wt% NorHA | 2178.7 ± 127.2 | 73.1 ± 3.9 |
| Reference 23 | 8,000 | 82-86 | 6 wt% PEGDM | 240 ± 84 | 12.5 |
| Reference 23 | 6,000 | 82-86 | 6 wt% PEGDM | 270 ± 87 | 9 |
| Reference 23 | 4,600 | 82-86 | 6 wt% PEGDM | 630 ± 79 | 8.2 |
| Reference 27 | 2,000,000 | 5 | 1 wt% GMHA | 109.4 ± 16.3* | 644 |
| Reference 27 | 2,000,000 | 7 | 1 wt% GMHA | 131.3 ± 6.4* | 619 |
| Reference 27 | 2,000,000 | 11 | 1 wt% GMHA | 154.5 ± 28.2* | 539 |

### Effects of Hydrogel Mechanical Properties on OPC Morphology, Proliferation, and Metabolic Activity

After synthesizing hydrogels with mechanical properties resembling CNS tissue, we encapsulated OPCs in NorHA hydrogels and assessed their viability through confocal fluorescence microscopy over 7 days of culture. Live/Dead imaging showed that OPCs could remain viable in all hydrogel formulations (**Figure 3**). Since NorHA hydrogels have larger mesh sizes than PEGDM hydrogels, we would also expect OPC spheroid volumes to be larger. Representative max projections of OPCs in hydrogels (**Figure 3**) visually show similarities between the 1 and 1.5 wt% groups over the 7 day period in comparison to the 2 wt% hydrogels. The vast majority of cells in these hydrogels exhibited rounded morphologies, which corroborates previous work showing that OPC process extensions inside 3D hydrogels decrease significantly when the elastic modulus is > 120 Pa.^31^

**Figure 3:**
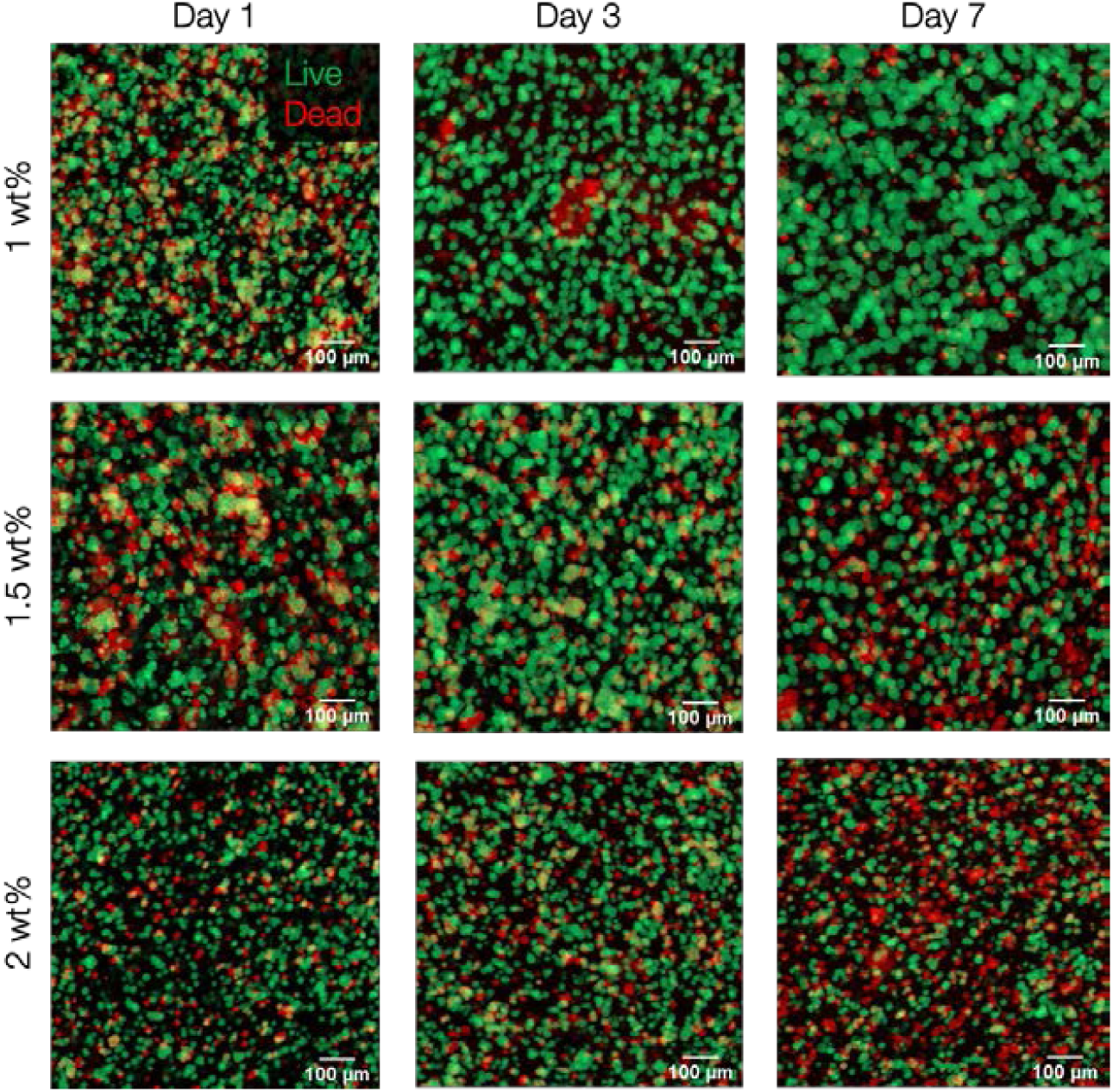
OPC viability in NorHA hydrogels. Standard deviation z-stack projections (770-1930 µm depth) of GFP+ OPCs (live cells, *green*) and ethidium homodimer-labeled dead cells (*red*) in NorHA hydrogels. 1 wt% hydrogels appear to show the largest average spheroid volumes and highest viability after 7 days of culture.

We used confocal live/dead images to measure OPC spheroid size (**Figure 4**). We previously reported an average OPC spheroid diameter of 21 µm for ∼ 600 Pa stiffness and 10 nm mesh size PEGDM hydrogels.^23^ In the current study, average OPC spheroid diameters were at least 1.5 fold greater in hydrogels of ∼ 795 Pa stiffness and 58 nm mesh size. NorHA hydrogels of ∼ 170 Pa and 67 nm mesh size had 2-2.5 fold greater spheroid diameters than the ∼ 600 Pa stiffness and 10 nm mesh size PEGDM hydrogels. This is consistent with the concept that larger mesh sizes lead to larger spheroid sizes, especially in hydrogels that are not engineered to degrade via hydrolysis or cell-secreted enzymes. In terms of absolute spheroid size, lower stiffness 1 wt% hydrogels did trend toward supporting the largest spheroid volumes after a 7 day culture period (**Figure 4)**.

**Figure 4:**
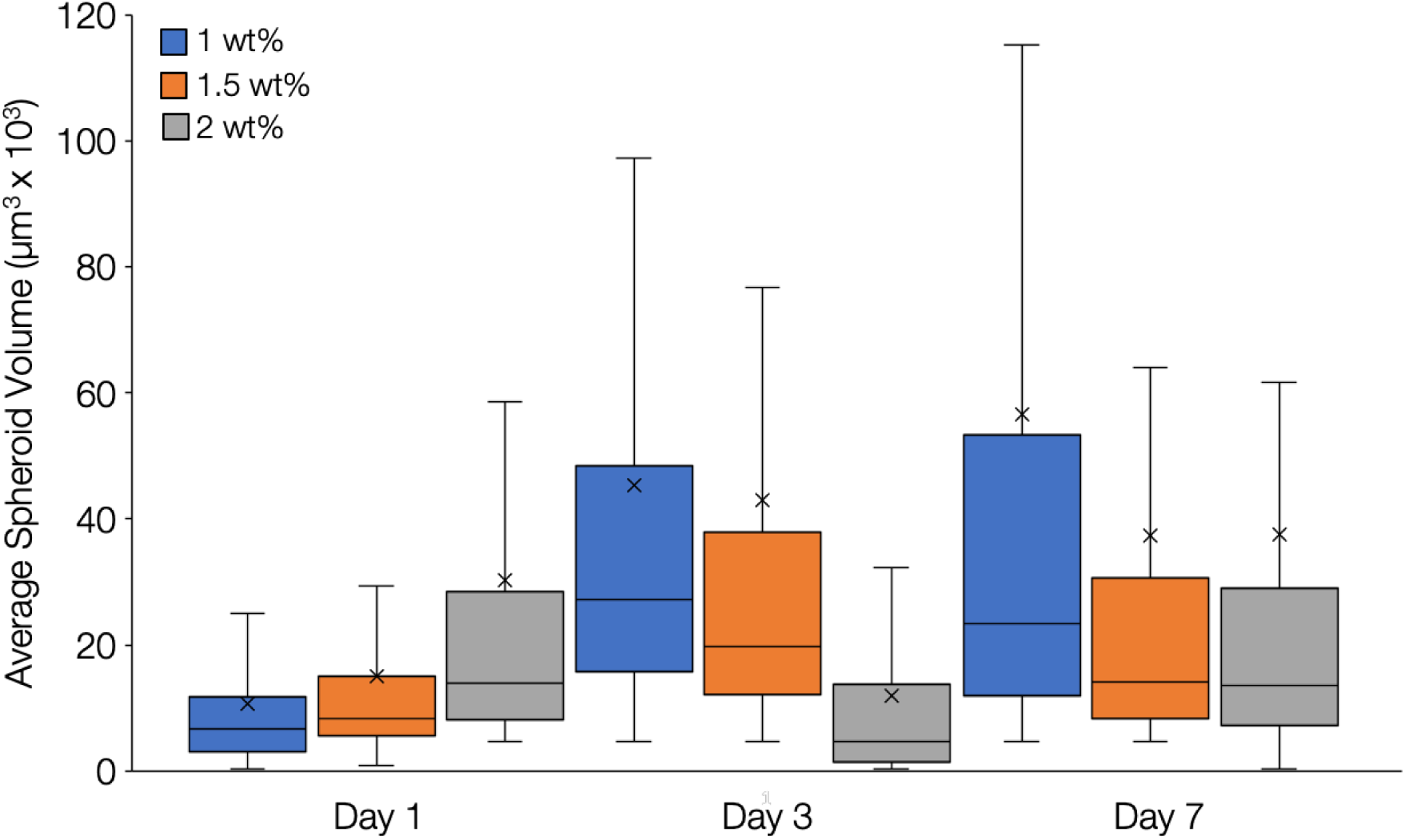
OPC spheroid volume. Average live spheroid volume was determined using the 3D object counter tool in ImageJ. 1 wt% hydrogels supported the largest average spheroid volume after 7 days of culture. Data presented for each group includes the mean (*x*) and median (*bar*). Whiskers represent the 1^st^ and 4^th^ quartiles while boxes represent the 2^nd^ and 3^rd^ quartiles. *n* = 15 (1 wt%) or 12 (1.5, 2 wt%) hydrogels per experimental group. At least 3500 spheroids were analyzed per experimental condition.

When normalized to hydrogel volume to consider differences in swelling, average spheroid volumes of 1 wt% and 1.5 wt% NorHA hydrogels were equivalent **(Figure S2)**. It was found that both hydrogel weight percentage and day of culture influenced the normalized average spheroid volume with spheroids in 1 wt% and 1.5 wt% hydrogels showing significantly higher levels than spheroids in 2 wt% hydrogels (α < 0.05). Normalized average spheroid volume was also significantly increased for all groups at days 3 and 7 compared to day 1 (α < 0.05). We hypothesized that our hydrogel groups could support different numbers of spheroids, which would change the total spheroid volume compared to average spheroid measurements. The time of culture affected the total spheroid volume with significantly higher levels quantified at day 7 compared to day 1 (α < 0.05) (**Figure S3**).

We next quantified OPC viability by measuring the total live spheroid volume to dead spheroid volume ratio (**Figure 5**) and compared our results to previous 3D OPC culture studies. Egawa et al. reported viability of OPCs in collagen/hyaluronan hydrogels, supported in part through lactate dehydrogenase (LDH) release, with values peaking at around 75%.^32^ In our NorHA hydrogels we used the total live to dead volume ratio to report viability with ratios of 6.7, 4.2, and 2.1 quantified at day 7 for the 1, 1.5, and 2 wt% hydrogel groups respectively. These ratios correspond to 87%, 81%, and 73% viability for the 1, 1.5, and 2 wt% hydrogel groups respectively.

**Figure 5:**
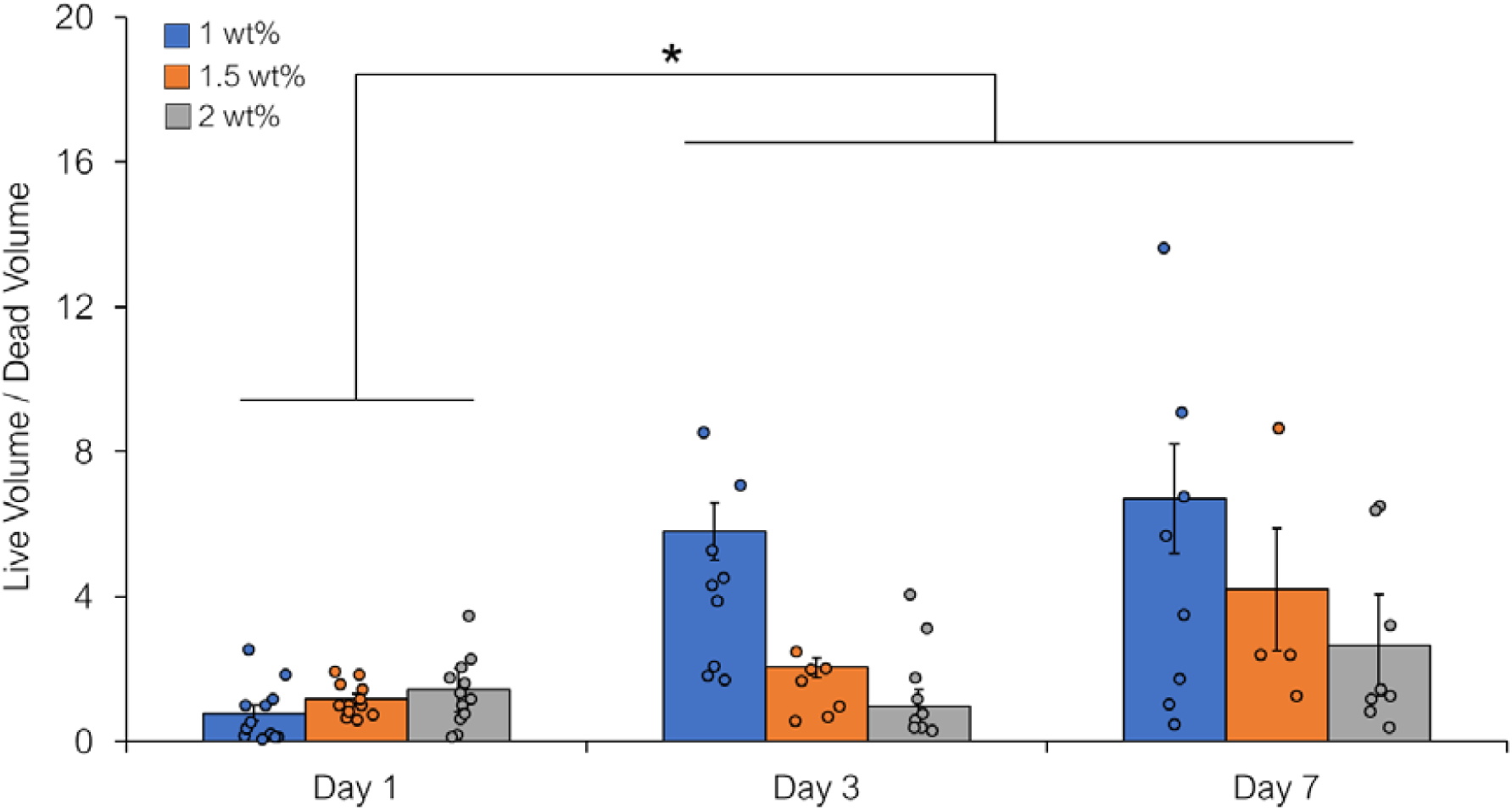
OPC live/dead volume ratio. The ratio of total live spheroid volume to dead spheroid volume was calculated for all experimental groups with 1 wt% hydrogels supporting higher live volume ratios compared to other groups at days 3 and 7 (α < 0.05). For each experimental group, bar height represents the average live/dead volume ratio, circles represent individual data points, and error bars represent the standard error of the mean. * indicates a statistically significant difference between day 1 compared to days 3 and 7 (α < 0.05). *n* = 4-13 hydrogels per experimental group.

After measuring OPC viability and spheroid formulation, we next sought to track OPC metabolic health in HA hydrogels over a 7 day culture period. We first measured OPC mitochondrial metabolic activity by measuring AlamarBlue reduction (**Figure 6**). This assay can be used to assess OPC metabolic activity in hydrogel samples over an extended period of time in a non-destructive manner by introducing a resazurin dye that is reduced to the fluorescent byproduct resorufin in the electron transport chain without interfering with it.^33^ The dye is reduced before oxidative phosphorylation, and requires 2H^+^ and 2e^-^ from the Krebs cycle per molecule to be reduced. We found that metabolic activity levels were similar amongst hydrogel groups at each time point with significantly higher values measured at day 7 compared to days 1 and 3 for all groups (α < 0.05).

**Figure 6:**
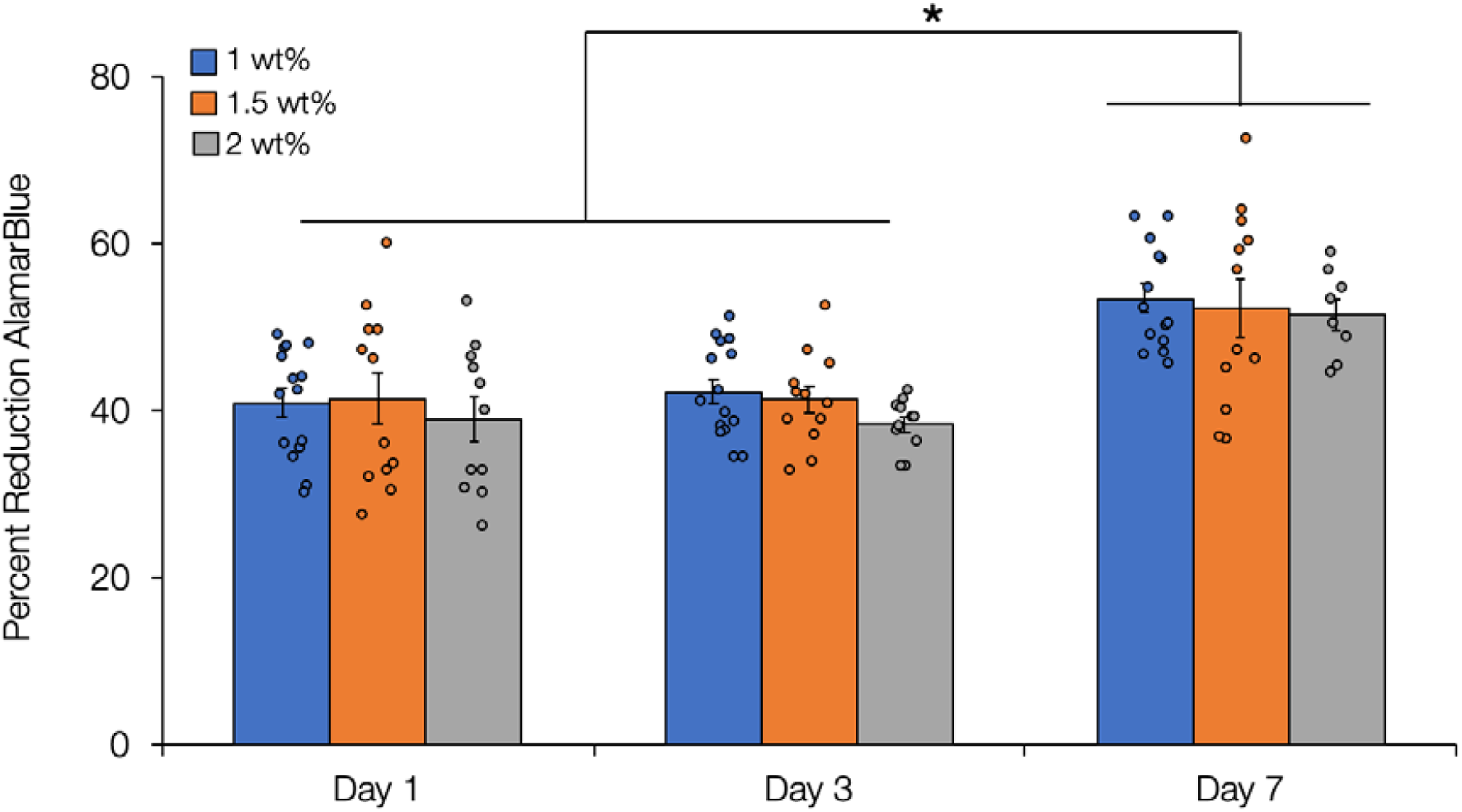
OPC mitochondrial metabolic activity. OPC metabolic activity in HA hydrogels was measured by AlamarBlue reduction. Hydrogels supported similar levels of metabolic activity at each time point with higher values measured at day 7. For each experimental group, bar height represents the average percentage reduction of AlamarBlue, circles represent individual data points, and error bars represent the standard error of the mean. * indicates a statistically significant increase in metabolic activity at day 7 compared to days 1 and 3 (α < 0.05). *n* = 9 (1 and 2 wt%) or 12 (1.5 wt%) hydrogels per experimental group.

We also measured OPC DNA concentration in hydrogels and observed similar trends to those found with AlamarBlue metabolic activity measurements with significantly higher DNA levels quantified at day 7 compared to days 1 and 3 for all hydrogel groups (**Figure 7A**). This increase in DNA content indicates that OPCs are undergoing substantial proliferation inside the NorHA hydrogels. These results are comparable to our previous findings with PEGDM hydrogels, where a statistically significant difference in DNA concentration was found between 240 Pa and higher stiffness (560, 1910 Pa) hydrogels.^23^

**Figure 7:**
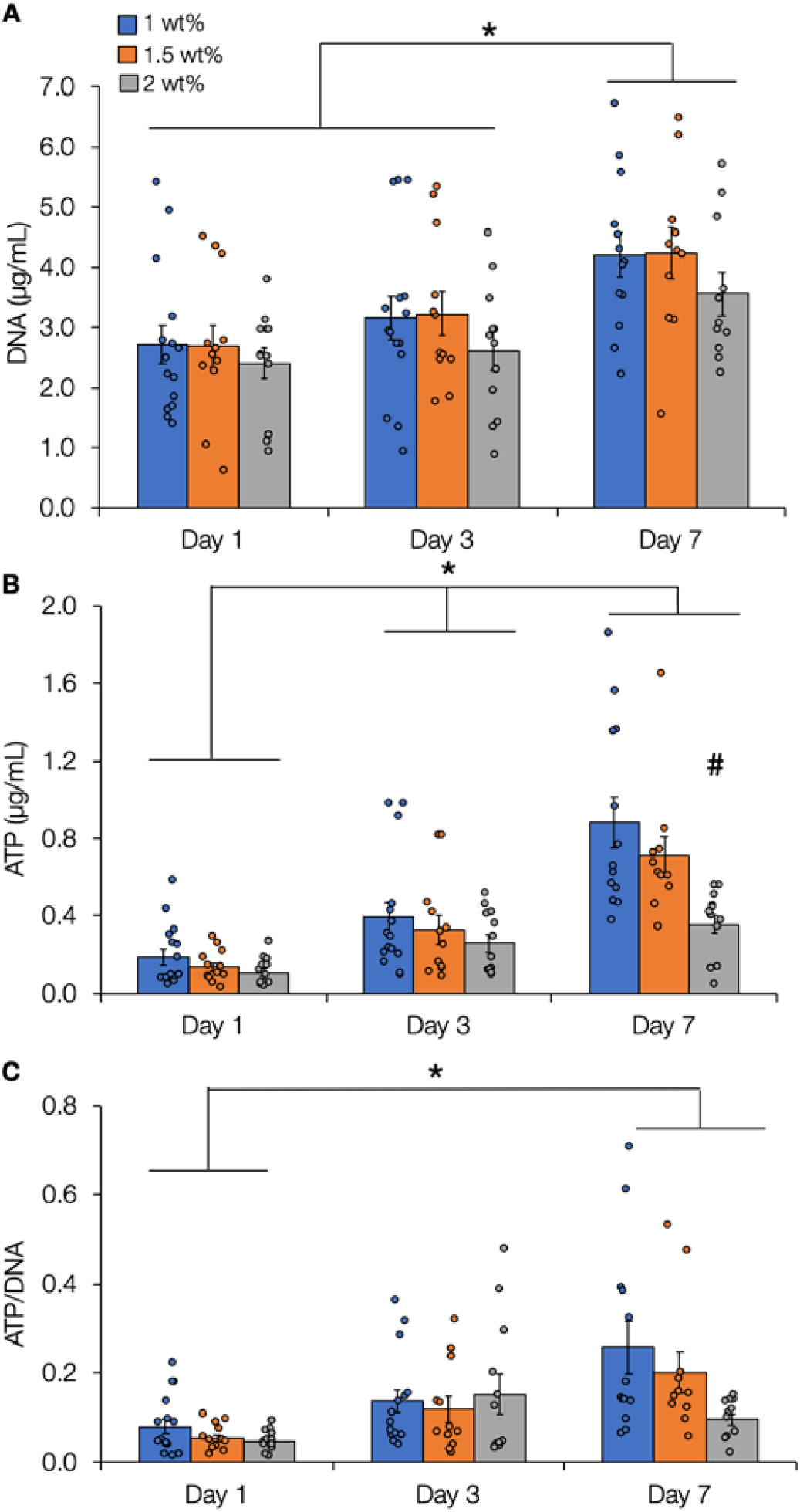
OPC ATP and DNA levels. (A) OPCs encapsulated in 1, 1.5, or 2 wt% NorHA hydrogels all proliferated over a period of 7 days. (B) OPCs also showed progressively increasing ATP concentration over a period of 7 days for all hydrogel groups. (C) Normalizing ATP to DNA concentrations demonstrated higher metabolic activity per cell at day 7 for all hydrogel groups with a trend toward increasing levels for OPCs in lower stiffness hydrogels. For each experimental group, bar height represents the average value for the data set, circles represent individual data points, and error bars represent the standard error of the mean. * indicates statistically significant differences between time points (α < 0.05), # indicates statistically significant differences between the 2 wt% and other hydrogel groups at a specific time point. *n* = 11-15 hydrogels per experimental group.

ATP concentration was also quantified as another measurement of OPC metabolic activity in HA hydrogels. ATP concentrations progressively increased in each hydrogel group as a function of culture time (**Figure 7B**). The lowest stiffness hydrogels demonstrated the highest ATP concentrations with significantly lower ATP levels measured in the stiffest (2 wt%) hydrogels at day 7. This corroborates our previous work with PEGDM hydrogels where the stiffer hydrogels showed lower ATP activity.^23^ After normalization of ATP to DNA levels as a measure of average metabolic activity per cell (**Figure 7C**), it was found that the day of culture had a significant influence with higher levels observed at day 7 compared to day 1 for all hydrogel groups. Additionally a trend toward higher ATP per DNA for lower stiffness hydrogels was observed at day 7.

The two cell metabolic activity assays performed (AlamarBlue, Cell Glo) offer an interesting point of comparison in our assessment of overall OPC health in HA hydrogels. The Cell Glo assay only monitors ATP production, including the 2 molecules of ATP produced by glycolysis and 30 molecules produced through oxidative phosphorylation per glucose molecule in cellular respiration.^34^ In contrast, the AlamarBlue assay measures NADH-dependent enzymatic reduction of resazurin to the fluorescent byproduct resorufin. The mechanistic differences between these assays suggests that Cell Glo may highlight sharper, more specific differences in energy metabolism between OPCs in different hydrogels, which is indeed what we observed (**Figure 6, 7)**.

Although our ATP/DNA results suggest that there is an inverse relationship between metabolic activity per cell and hydrogel stiffness, this relationship is non-linear. We also observe a non-linear relationship between hydrogel stiffness and mesh size. A 4.6-fold increase in stiffness (1.5 wt% in comparison to 1 wt%) corresponds to a ∼ 11.6% reduction in mesh size (**Table 1**). A 12-fold increase in stiffness (2 wt% in comparison to 1 wt%) corresponds to a ∼ 14.5% decrease in mesh size (**Table 1**). This is similar to our previous work with PEGDM hydrogels where a 2.6- fold increase in stiffness led to a mesh size change from 9.47 nm to 6.61 nm (30% decrease) as well as significant decreases in ATP concentration.^23^ The evidence from this study and our previous work suggests that OPC metabolic activity per cell is affected by the non-linear relationship between hydrogel stiffness and mesh size. Considering the hydrodynamic radii of most proteins is on the order of 1-6 nm,^35^ all hydrogels could likely support adequate diffusion of necessary biomolecules. Interestingly, 1.5 wt% and 2 wt% hydrogels supported similar average OPC spheroid volumes (**Figure 4**) despite reduced metabolic activity per cell measurements in 2 wt% hydrogels. However, normalizing the average spheroid volumes to the hydrogel volume (**Figure S2**) leads to trends that correlate more closely to metabolic activity results (larger spheroids correlate to higher levels of metabolic activity), suggesting that quantification of spheroid volumes in a way that accounts for differences in hydrogel swelling helps explain our metabolic activity results. These results provide an important point of comparison for material systems and strongly indicate that stiffness should not be the sole criteria of interest in 3D mechanobiological studies, at least for OPCs. It should also be noted that the material system explored here does not give rise to substantial remodeling in the 7 day time course of our experimental study, whereas remodeling may be an important component in both maintenance^36^ and differentiation^37^ of neural lineages.

## Conclusions

We successfully synthesized and characterized NorHA hydrogels (1, 1.5, and 2 wt%) with mechanical properties resembling CNS tissue (storage moduli of ∼ 170, 790, and 2180 Pa respectively). Encapsulated OPCs showed good viability in all HA hydrogel formulations over 7 days of culture. OPCs formed progressively larger spheroids over the culture time with the largest spheroids formed in the lower stiffness (1 wt%) hydrogels with the largest mesh sizes. Average live spheroid volumes were greater than those previously observed in PEGDM hydrogels, perhaps due to larger mesh sizes in the HA hydrogels. Finally, we showed that HA hydrogels supported progressively increasing metabolic activity levels as measured by both AlamarBlue (mitochondrial metabolic activity) and Cell Glo (ATP) assays where ATP levels normalized to DNA were higher in more compliant HA hydrogels. Our results suggest that 3D HA hydrogels could be useful for the growth and expansion of OPCs, and that improved OPC metabolic health is favored in more compliant, larger mesh size, hydrogels.

## Acknowledgments

We gratefully acknowledge financial support from NSF grant 1904198 and the Translational Health Research Institute of Virginia for a career development award to KJL. We also acknowledge the Keck Center for Cellular Imaging at the University of Virginia for the usage of the Zeiss 510 microscopy system, NIH grant S10 RR021202 01. Finally we would like to thank the University of Virginia for awarding the Fogg Fellowship (DBU).

