## Supplemental information for "3D Hyaluronic Acid Hydrogels for Modeling Oligodendrocyte Progenitor Cell Behavior as a Function of Matrix Stiffness"

[*]Co-corresponding authors

#### Supplemental Information

Additional equations for determining crosslinking density and mesh size

$$W_{f,r} = \frac{m_d}{m_{w,r}} \qquad \text{(S1)}$$

$m_d = dry\ mass$

$m_{w,r} = wet\ mass\ of\ relaxed\ hydrogel\ (right\ after\ curing)$

$W_{f,r} = relaxed\ hydrogel\ dry\ mass\ weight\ fraction$

$$W_{f,s} = \frac{m_d}{m_{w,s}} \qquad \text{(S2)}$$

$m_{w,s} = wet\ mass\ of\ swollen\ hydrogel$

$W_{f,s} = swollen\ hydrogel\ dry\ mass\ weight\ fraction$

$$v_{2,r} = \left[1 + \frac{\left(\frac{1}{W_{f,r}}\right)\rho_{HA}}{\rho_{H2O}}\right]^{-1} \qquad \text{(S3)}$$

$\rho_{HA} = density\ of\ dry\ hyaluronic\ acid\ polymer$

$\rho_{H2O} = density\ of\ solvent\ (water)$

$$v_{2,s} = \left[1 + \frac{\left(\frac{1}{W_{f,s}}\right)\rho_{HA}}{\rho_{H2O}}\right]^{-1} \qquad \text{(S4)}$$

### Supplementary Figures

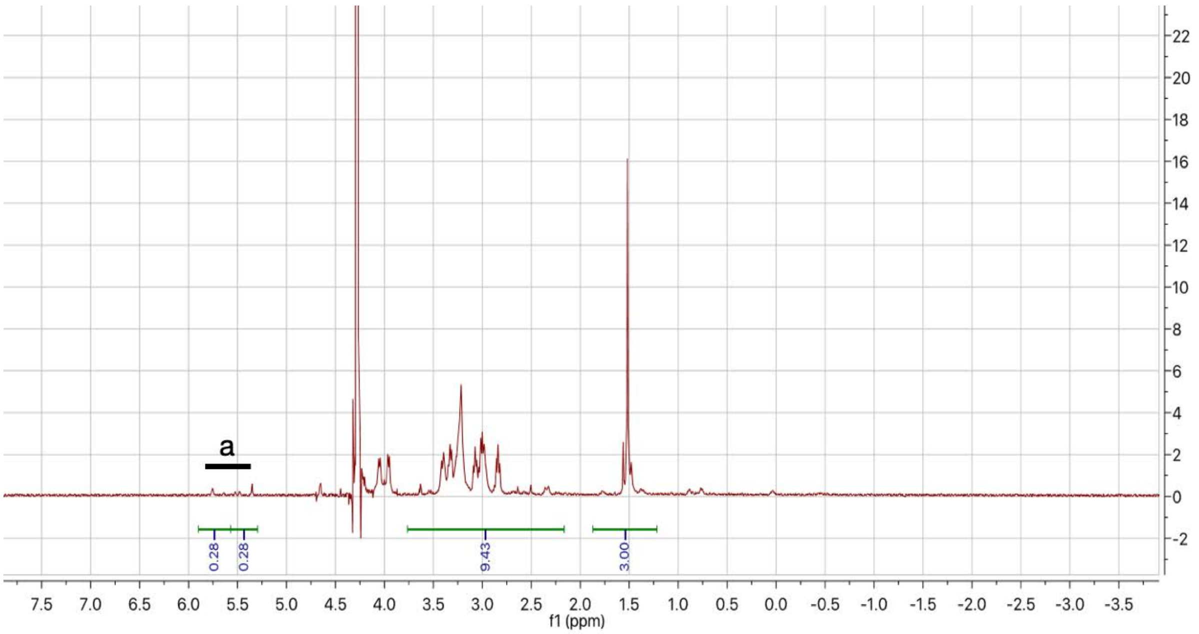


**Figure S1: $^1$H NMR spectrum of norbornene-modified hyaluronic acid (NorHA).** The methyl peak at 1.5 ppm is used as an integration standard to determine the degree of norbornene functionalization from the average integrations of the *endo*/*exo* vinyl protons at 5.2 and 5.7 ppm (a). The degree of modification of carboxylic acid groups on HA was determined to be 28%.

| Sample Type | Mass Swelling Ratio (0 h) | Mass Swelling Ratio (24 h) | Mass Swelling Ratio (48 h) | Volumetric Swelling Ratio (0 h) | Volumetric Swelling Ratio (24 h) | Volumetric Swelling Ratio (48 h) | % Change in Swelling 0 to 24 h | % Change in Swelling 24 to 48 h | Volume (0 h, $mm^3$) | Volume (24 h, $mm^3$) | Volume (48 h, $mm^3$) |
|---|---|---|---|---|---|---|---|---|---|---|---|
| 1 wt% NorHA | 54.4 ± 7.4 | 59.7 ± 4.9 | 57.2 ± 3.6 | 66.6 ± 9.0 | 73.1 ± 6.0 | 70.1 ± 4.5 | 9.7 ± 6.1 | - (4.5 ± 2.7) | 272.5 ± 19.6 | 335.3 ± 24.4 | 300.0 ± 44.1 |
| 1.5 wt% NorHA | 44.7 ± 4.6 | 52.2 ± 15.8 | 47.0 ± 1.5 | 54.7 ± 5.7 | 63.9 ± 19.4 | 57.5 ± 1.8 | 24.9 ± 26.9 | - (6.0 ± 2.0) | 247.9 ± 67.1 | 281.2 ± 52.9 | 275.4 ± 65.7 |
| 2 wt% NorHA | 38.0 ± 4.2 | 42.9 ± 2.6 | 43.0 ± 0.8 | 46.5 ± 5.1 | 52.5 ± 3.2 | 52.7 ± 1.0 | 27.0 ± 3.5 | - (2.4 ± 1.2) | 276.5 ± 9.7 | 365.4 ± 12.5 | 325.3 ± 17.6 |

**Table S1: Hydrogel swelling ratios.** NorHA hydrogels show moderate levels of swelling after 24 h with larger swelling ratios measured for lower weight percentage, lower stiffness hydrogels. Values are averages +/- standard deviation for 3 samples at each time point.

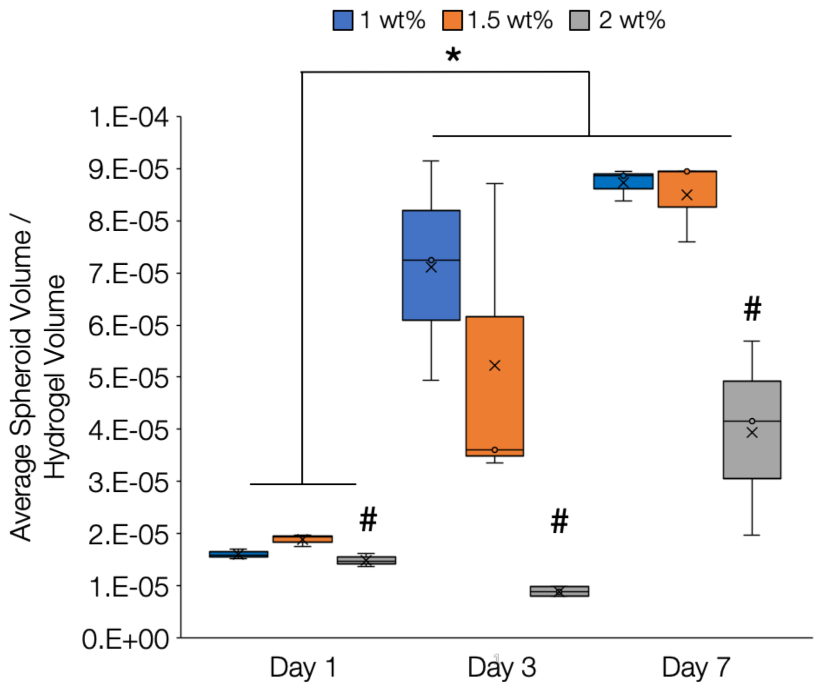


**Figure S2: Average OPC spheroid volume normalized to hydrogel volume.** The average volume of each live cell spheroid was determined using the 3D object counter tool in ImageJ and normalized to hydrogel volume. Normalized average spheroid volume was significantly increased at days 3 and 7 compared to day 1 with significantly lower values measured for 2 wt% hydrogels at all time points. Data presented for each group include the mean (*x*) and median (*bar*). Whiskers represent the 1$^{st}$ and 4$^{th}$ quartiles while boxes represent the 2$^{nd}$ and 3$^{rd}$ quartiles. * indicates statistically significant differences between time points, # indicates statistically significant differences between the 2 wt% and other hydrogel groups at a specific time point. $n$ = 15 (1 wt%) or 12 (1.5, 2 wt%) hydrogels per experimental group. At least 3500 spheroids were analyzed per experimental condition.

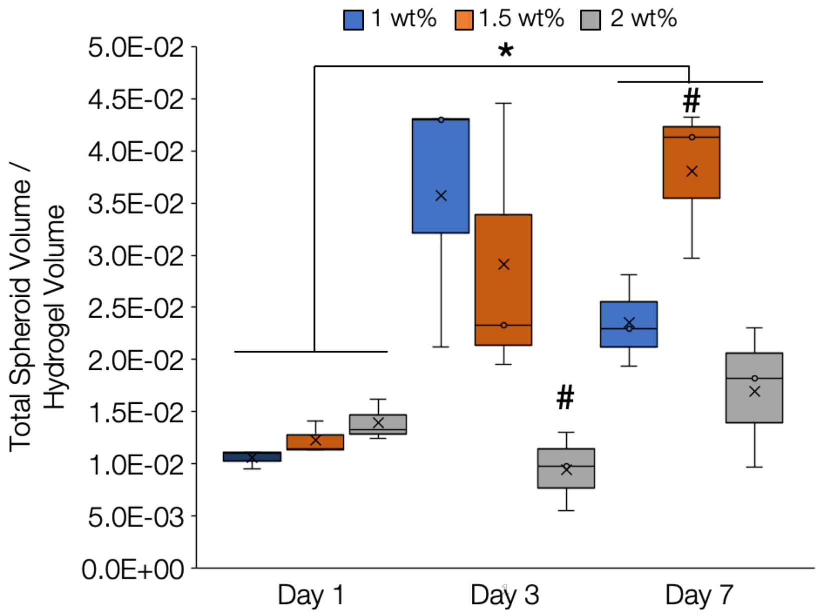


**Figure S3: Total OPC spheroid volume normalized to hydrogel volume.** The total volume of live cell spheroids (summation) was determined using the 3D object counter tool in ImageJ and normalized to hydrogel volume. Normalized total spheroid volume was significantly increased at day 7 compared to day 1. Significantly lower values were measured for 2 wt% hydrogels at day 3 while significantly higher values were measured for 1.5 wt% hydrogels at day 7. Data presented for each group include the mean (*x*) and median (*bar*). Whiskers represent the 1st and 4th quartiles while boxes represent the 2nd and 3rd quartiles. * indicates statistically significant differences between time points, # indicates statistically significant differences from other hydrogel groups at a specific time point. *n* = 15 (1 wt%) or 12 (1.5, 2 wt%) hydrogels per experimental group. At least 3500 spheroids were analyzed per experimental condition.
